# Unsupervised domain classification of AlphaFold2-predicted protein structures

**DOI:** 10.1101/2024.08.21.608992

**Authors:** Federico Barone, Alessandro Laio, Marco Punta, Stefano Cozzini, Alessio Ansuini, Alberto Cazzaniga

**Affiliations:** Area Science Park, Research and Technology Institute, Trieste, 34149, Italy; University of Trieste, Trieste, 34127, Italy; SISSA, International School for Advanced Studies, Trieste, 34136, Italy; ICTP, International Centre for Theoretical Physics, Trieste, 34151, Italy; IRCCS San Raffaele Institute, Center for Omics Sciences, Milan, 20132, Italy; IRCCS San Raffaele Institute, Unit of Immunogenetics, Leukemia Genomics and Immunobiology, Division of Immunology, Transplantation and Infectious Disease, Milan, 20132, Italy

## Abstract

The release of the AlphaFold database, which contains 214 million predicted protein structures, represents a major leap forward for proteomics and its applications. However, lack of comprehensive protein annotation limits its accessibility and usability. Here, we present DPCstruct, an unsupervised clustering algorithm designed to provide domain-level classification of protein structures. Using structural predictions from AlphaFold2 and comprehensive all-against-all local alignments from Foldseek, DPCstruct identifies and groups recurrent structural motifs into domain clusters. When applied to the Foldseek Cluster database, a representative set of proteins from the AlphaFoldDB, DPCstruct successfully recovers the majority of protein folds catalogued in established databases such as SCOP and CATH. Out of the 28,246 clusters identified by DPCstruct, 24% have no structural or sequence similarity to known protein families. Supported by a modular and efficient implementation, classifying 15 million entries in less than 48 hours, DPCstruct is well suited for large-scale proteomics and metagenomics applications. It also facilitates the rapid incorporation of updates from the latest structural prediction tools, ensuring that the classification remains up-to-date. The DPCstruct pipeline and associated database are freely available in a dedicated repository, enhancing the navigation of the AlphaFoldDB through domain annotations and enabling rapid classification of other protein datasets.

## Introduction

The recent release of the AlphaFold database (AlphaFoldDB) [1, 2], containing predictions for nearly 214 million protein structures, marked a significant breakthrough in proteomics, with applications ranging from structure-based drug discovery to variant effect prediction [3, 4]. With *in silico* structural information now available at a scale comparable to sequence data, the potential for new insights into protein evolution and function has dramatically expanded. However, despite the success of these structural predictions, lack of comprehensive protein annotation constitutes a significant obstacle toward taking full advantage of this vast dataset.

A common approach for leveraging such large and redundant databases is using clustering techniques, which partition data according to diverse criteria. These methods have been successfully applied to large protein sequence databases, where they have been used to formulate functional hypotheses based on homology, complementing and extending human curated annotations contained in datasets such as Pfam [5]. This annotation process can be made fully automatic by exploiting unsupervised clustering techniques: for instance, in previous work we successfully classified protein domains based on sequence similarity, identifying 45,000 protein clusters in the UniRef50 database [6]. This approach provided valuable insights into protein family organization, hinting at occasional misclassifications and helping to uncover previously unknown families, some of which are now part of the Pfam 36.0 release.

Sequence-based classification methods have inherent limitations, primarily due to the difficulty in finding distant homologs, resulting in large portions of current protein databases lacking annotation [7, 8]. Leveraging structural data can facilitate the identification of remote homology relationships (i.e., between proteins with less than 20% sequence similarity), thereby improving coverage. In addition, structure often correlates more closely with protein function, aiding in the development of functional hypotheses. For these reasons, over the past decade, several initiatives have focused on classifying proteins based on structural and evolutionary relationships, using the PDB database as a primary source of information [9, 10, 11, 12, 13]. Extending these efforts to the scale of AlphaFoldDB could enable a more accurate and comprehensive classification of the protein universe.

Significant progress has already been made in this direction. Durairaj et al. combined a sequence-based protein similarity network with AlphaFold structural predictions to identify new folds and extend the Pfam database [14]. Barrio-Hernandez et al. presented the Foldseek Cluster database, built by clustering the AlphaFoldDB database at 50% sequence similarity and 90% structural overlap thresholds. The resulting database contains 15 million clusters and their associated representative proteins [15, 16]. Their work focused on full-protein clusters, but they also discussed the importance of domain-level classification, for which they proposed a proof-of-concept algorithm. Domain-level classification is particularly valuable because proteins can be divided into functional modules, and these regions tend to be better conserved evolutionarily. An important contribution in this regard is the work of Bordin et al., in which the authors used deep learning techniques to automatically identify and assign protein domains from AlphaFold models to the CATH classification [17, 18]. While their approach achieved impressive coverage, it was designed to fit the CATH classification criteria, which differs from other classification schemes such as the ones of SCOP[10]. It is also computationally demanding, requiring several months on a computing cluster to generate the classification [18]. This high computational cost can be problematic when dealing with larger datasets, such as those from metagenomics, or with the frequent updates to predicted structure databases.

In this study, we classify protein domain families of AlphaFoldDB with DPCstruct, an enhanced implementation of the DPCfam algorithm aimed at reliably identifying structural domains [6]. Leveraging structural predictions from AlphaFold2 and all-against-all local alignments performed with Foldseek, DPCstruct singles out recurrent structural motifs and groups them into clusters of protein domains. Unlike previous methods, our approach is entirely unsupervised and does not depend on prior knowledge of protein domains for classification, making it ideal for uncovering relationships in uncharted regions of the protein universe and for future applications in metagenomics [19, 20].

DPCstruct required only 48 hours of computational time to recover the majority of protein folds catalogued in well-established structural databases such as SCOP and CATH, which are the result of decades of experimentation combined with extensive manual curation [10, 11]. Our results also show strong agreement with Pfam clan-level annotation and capture remote homologs’ relationships between Pfam families. Additionally, DPCstruct identifies numerous novel protein clusters that may represent new protein families or folds, further contributing to the annotation and functional prediction of the vast uncharacterized protein space.

In summary, by leveraging the comprehensive structural predictions from AlphaFoldDB and the efficient alignment capabilities of Foldseek, DPCstruct provides an innovative and scalable solution for the structural identification and classification of protein domains.

## Results

### DPCstruct in numbers

In this work, we apply DPCstruct to the 15 million representative structures from the Foldseek Clusters database [16]. Representative structures in Foldseek Clusters are obtained by reducing redundancy in AlphaFoldDB under the requirement that any two proteins share no more than 50% sequence identity and 90% structural overlap (see Figure 1(a)). The goal of DPCstruct is finding *domains*, namely consecutive subsets of a protein sequence which form a structural modulus. Therefore, a Foldseek Cluster representative may contain more than one domain, and the same domain can be contained in many representatives.

**Fig. 1:**
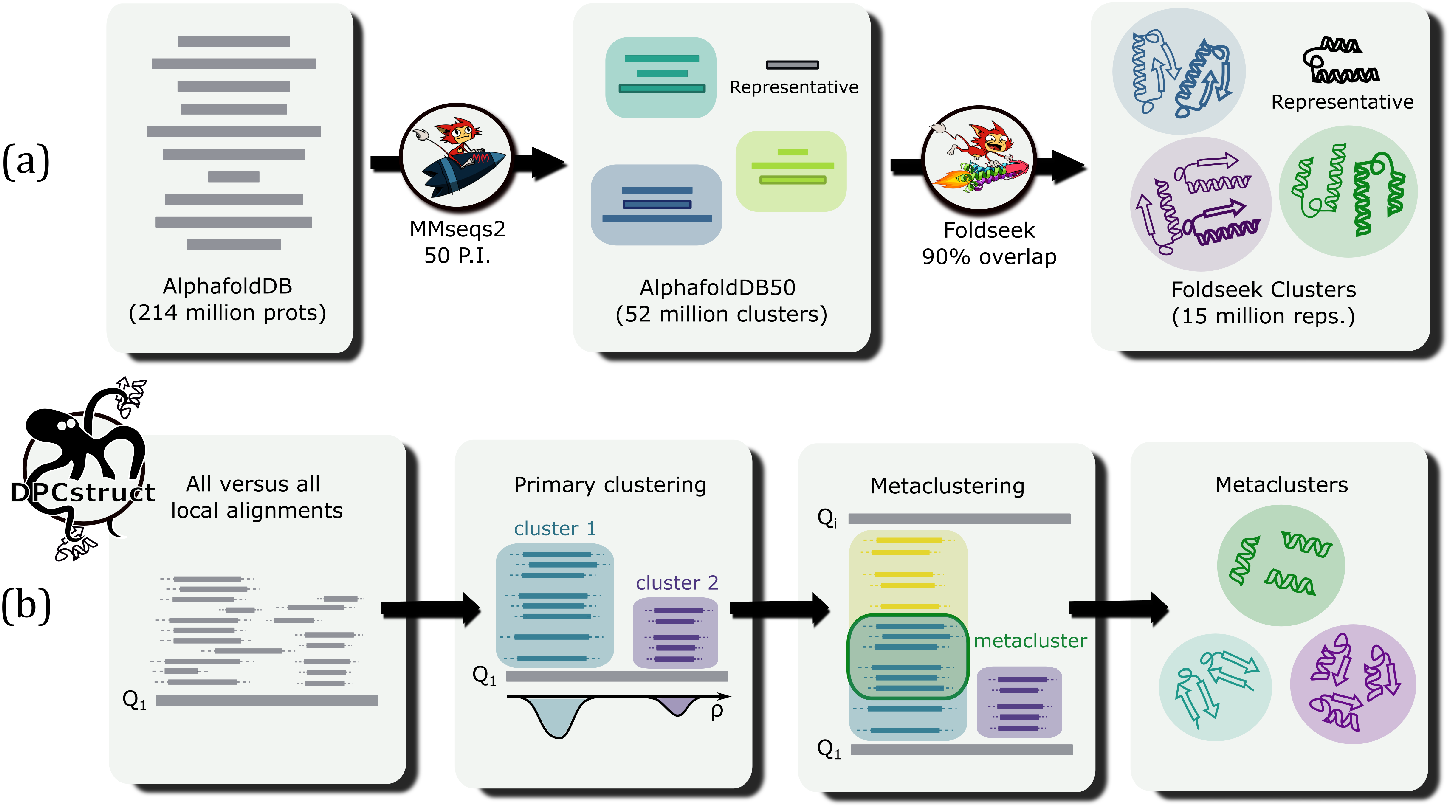
**(a)** The dataset used as input for DPCstruct is the Foldseek Clusters database without fragments, a full protein, non-redundant version of the AlphaFoldDB with approximately 15 million representatives [16] **(b)** The DPCstruct pipeline consists of three steps: First, an all-versus-all local alignment using Foldseek is performed on the input database. Second, performing density-based clustering for each query protein to identify clusters of local alignments. Finally, merging these clusters into metaclusters, each consisting of a list of non-redundant, structurally related protein domains.

The first step in our pipeline consists in performing an all-versus-all local alignment of the input data using the Foldseek structural aligner. For each query protein, we group local alignments extending over similar regions of the query into clusters of putative domains, or primary clusters (see Figure 1(b)). Of the 15 million query proteins, only 16% featured at least 10 local alignments to other proteins, the minimum required to qualify as a primary cluster. The generally low level of local structural similarity observed among the representatives can be attributed to either a high degree of structural uniqueness or to a low quality of their AlphaFold-generated structures (see Supplementary Figure 1(a)). As we will see, this does not affect the quality of the final results. Indeed, the primary clusters that were identified correspond to proteins that represent the more densely populated Foldseek Clusters, accounting for approximately 74% of all entries in the AlphaFold database. Moreover, they are among those with higher confidence scores (see Supplementary Figure 1(b)).

Following the identification of the primary clusters, we proceeded with a secondary clustering step to merge these groups into metaclusters. The final output consisted of 1,642,061 domains distributed over 1,252,179 proteins, organized into 28,246 metaclusters based on their structural similarity. Most metaclusters exhibit a uniform distribution of domain lengths, with only 9% of them showing a ratio between standard deviation and mean greater than 0.2. The within-metacluster average domain length ranges from 30 to 1000 amino acids, with 97% of metaclusters having an average domain length of less than 400 amino acids.

The distribution of the number of domains within a metacluster follows a power-law curve which is characteristic of scale-free networks typically observed in many biological processes [21] (see Figure 2(a), green curve). This distribution of domain sizes has been modeled and measured for protein families, superfamilies, and folds within a single genome [22]. We extend these results by showing that DPCstruct metaclusters, corresponding to structurally related protein regions, exhibit the same behavior even when considering a wide variety of genomes from all domains of life. A similar power-law behavior is also observed for families detected by the DPCfam algorithm on UniRef50 dataset clustering by sequence similarity (violet curve) [6].

**Fig. 2:**
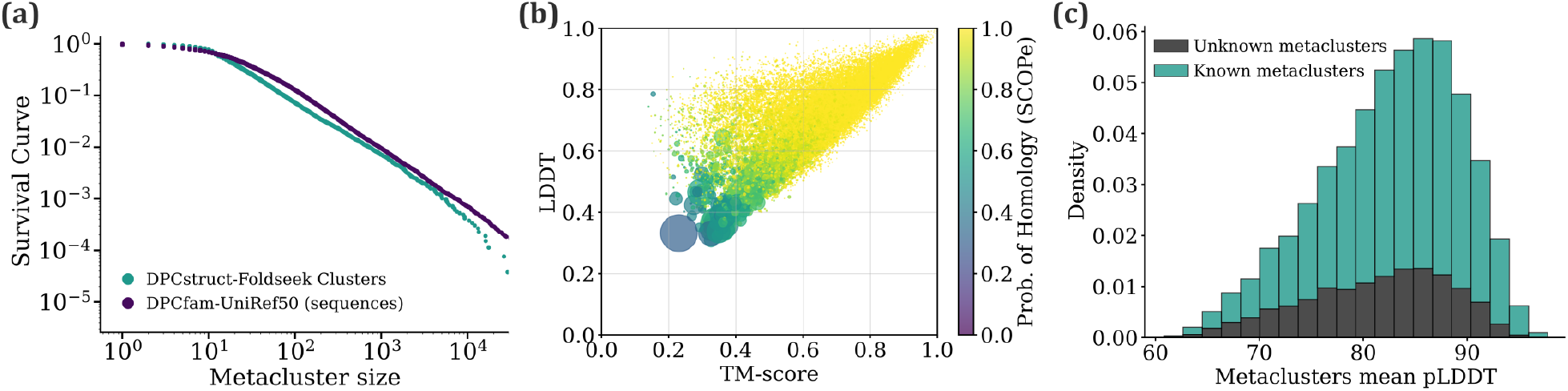
General properties of metaclusters. **(a)** Survival curve for metaclusters generated from Foldseek Clusters representatives structures (green) and, as reference, from UniRef50 sequences (violet) [6]. Both curves exhibit a power-law behaviour for over 3 order of magnitude. **(b)** Scatter plot showing the relationship between average LDDT and TM-score per metacluster. Marker size is proportional to metacluster size, and the color corresponds to the probability of homology according to SCOPe. All values are calculated using Foldseek. **(c)** Distribution of the mean predicted LDDT score per metacluster. Within each bin we show the proportion of metaclusters with and without annotations, labeled as known or unkown respectively.

To assess the structural consistency of domains assigned to the same class, we computed the average local distance difference test (LDDT) and template modelling (TM) score for each metacluster [23, 24]. These metrics were obtained by averaging all pairwise alignments of domain members within each metacluster. Results are summarized in Figure 2(b), where marker size is scaled proportionally to the metacluster size. Additionally, marker color represents the probability of homology according to SCOPe, introduced by Foldseek as an alternative method for estimating homology [15]. The median metacluster LDDT and TM-score values are 0.74 and 0.72, respectively. These values are in line with those observed for full protein clusters [16]. The majority of metaclusters (98%) exhibit an average LDDT and TM-score greater than or equal to 0.5, a commonly accepted threshold indicating structural similarity between proteins [25]. Only 38 metaclusters have LDDT and TM-score values lower than 0.4. Although these clusters are relatively few in number, they account for 14% of all the proteins for which a domain is detected by DPCstruct, within the Foldseek Clusters database. Manual inspection suggests that while these metaclusters could potentially be subdivided further, they still maintain a high level of structural similarity.

For each predicted structure, AlphaFold provides a predicted LDDT score (pLDDT), which is an estimate of the similarity between the prediction and the actual protein structure. To be considered in our pipeline, domains must have a pLDDT score higher or equal than 60%. In Figure 2(c), we show the distribution of mean pLDDT across DPCstruct metaclusters. Most metaclusters are composed of structures with high pLDDT scores, on average 82%, indicating high quality predictions.

### Coverage of reference structure-based domain databases

The capability of DPCstruct to retrieve protein families and folds was assessed by comparing our results with two established structure-based protein family databases, SCOP [10] and CATH [17]. Figure 3 shows the ratio of folds annotated by DPCstruct to non-annotated ones for different fold sizes, defined as the number of domains within each fold in the respective database. For a fold to be considered annotated by DPCstruct, we require that at least one domain in the fold is structurally aligned to a DPCstruct domain, with a TM-score or LDDT greater or equal than 0.5 and a fold sequence coverage of 0.8 or greater. The overall recall rate was 94% for SCOP folds and 86% for CATH. If we consider partially covered folds by reducing the coverage threshold to 0.5, total recall increases to 97% and 89%, respectively. As shown in Figure 3, the majority of non-annotated folds in both SCOP and CATH are small in size. DPCstruct is designed to identify putative protein families based on their recurrence in nature, which inherently makes the detection of rare folds more challenging. Upon closer examination of the larger non-annotated folds, we found that 9 out of the 10 largest CATH folds without DPCstruct annotation fall into one of two categories: those associated with viral proteins,which are not present in the AlphaFold Database [1] and consequently not part of the DPCstruct input dataset, and those labeled as irregular by the CATH architecture classification, which are proteins with anomalously low secondary structure content.

**Fig. 3:**
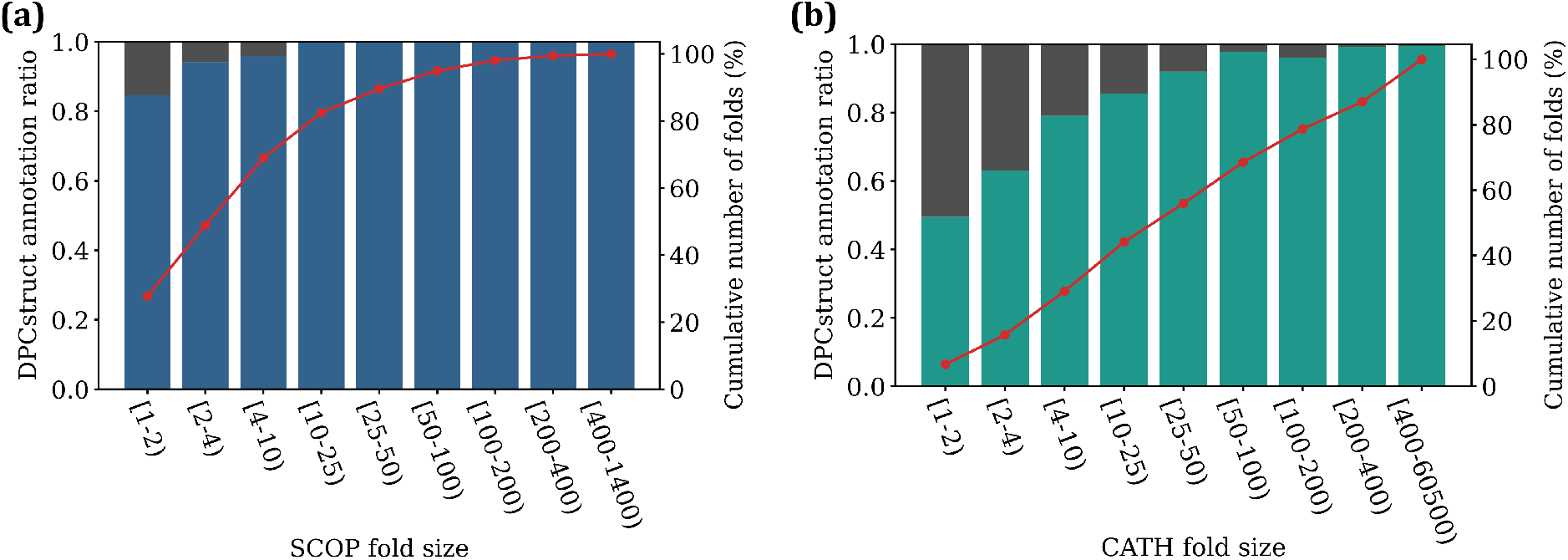
DPCstruct metaclusters coverage of folds as identified by reference structural domain databases (a) SCOP and (b) CATH. Each bin shows the proportion of annotated (blue or green) versus non annotated (gray) folds within a certain size, defined as the number of domains within the fold. For a fold to be covered, it must contain at least one domain covered by a DPCstruct domain, with a coverage threshold of 80% of the fold and a TM-score or LDDT greater than 0.5. The red curve represents the cumulative number of folds as a percentage of the total number in each database.

Note that there is no universally accepted definition of fold or family, even among the aforementioned reference databases. Nevertheless, by identifying recurring structural motifs, DPCstruct manages to retrieve almost all folds present in the reference databases, demonstrating its ability to capture known structural relationships. In the next sections we will see that DPCstruct has also potential to uncover novel relationships or folds.

### Consistency with respect to sequence-based Pfam database

To further validate our results, in particular their agreement with sequence-based annotations, we compared the labels assigned by DPCstruct with those from Pfam v36.0 [5]. Pfam is a widely used database that classifies protein domains based on known sequence similarities. Moreover, Pfam provides two hierarchical levels of classification: protein families, which aggregate closely related sequences, and clans, which group evolutionary related families with a typically lower degree of sequence similarity. Since DPCstruct identifies domains based on their structural similarity, which is effective for capturing distant homologs, we performed the comparison at the clan level. Pfam families without a clan were assumed to constitute their own clan.

For this comparison, we focused on metaclusters that have at least two structures with a corresponding Pfam annotation (14,423 in total). Among these, 70% exhibit a consistency score of 1, indicating that all structures contained the same Pfam annotation (see Methods). Conversely, Pfam clans have lower consistency scores, with only 38% of clans displaying perfect agreement with DPCstruct annotations (see Figure 4(a)). This lower consistency is largely due to Pfam predominantly identifying single-domain families, whereas DPCstruct metaclusters can include either single or multiple domain architectures, sometimes nested. We also identified potential misclassifications within Pfam annotations. When restricting the analysis to Pfam seed domains, which are manually curated entries, consistency scores improved, with 81% of metaclusters and 61% of Pfam clans achieving a score of 1.

**Fig. 4:**
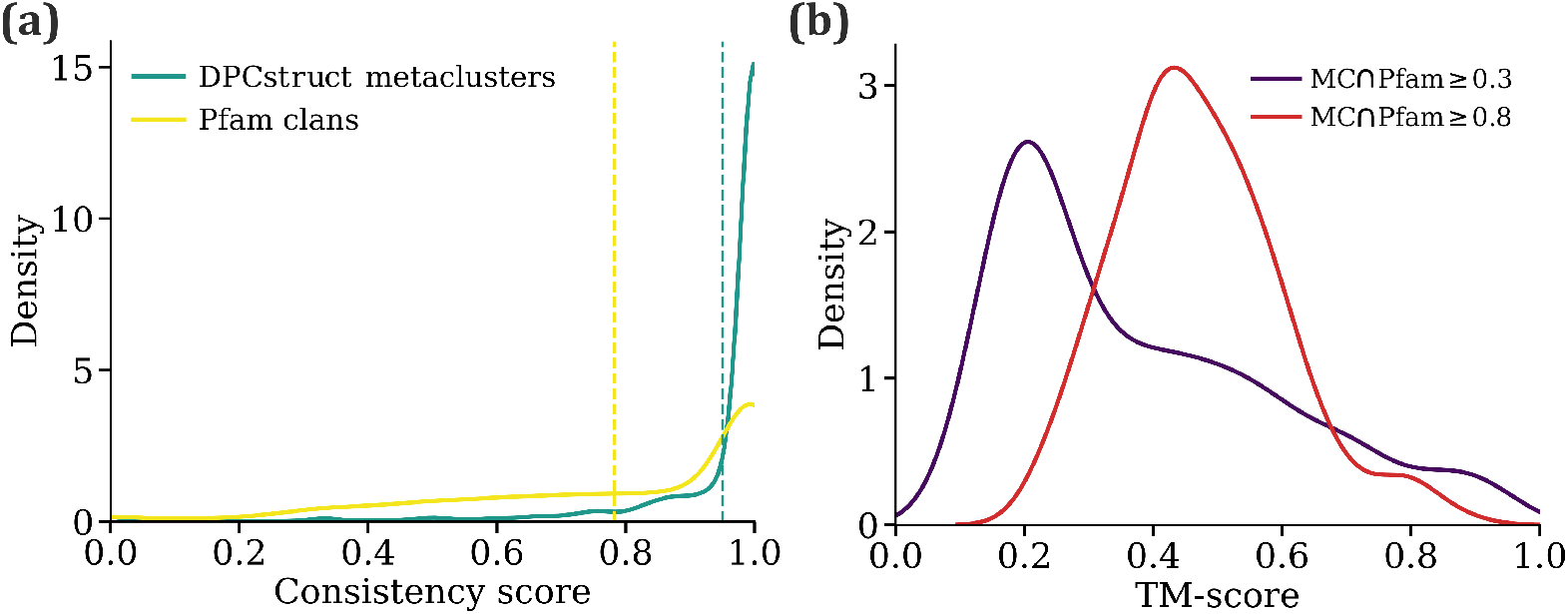
DPCstruct metaclusters and Pfam clans consistency. **(a)** Consistency score defined as the average number of equally labeled pairs of domains for DPCstruct classification with respect to Pfam clans (green) and Pfam clans with respect to DPCstruct metaclusters (yellow). Vertical lines indicate the mean of the distribution: 0.95 for DPCstruct and 0.78 for Pfam. **(b)** Average TM-score between domains of DPCstruct metaclusters containing two different Pfam labels. We consider Pfam labels with a minimum intersection with DPCstruct domains of 0.3 (violet) and 0.8 (red). The higher the intersection threshold the higher the average TM-score as we transition from metaclusters partially matching Pfam labels, generally part of architectures, to structurally aligned domains.

Metaclusters with a low consistency score are particularly interesting as they may reveal novel, previously unknown evolutionary relationships between clans. In what follows, we focused on the subset of metaclusters containing Pfam labels from two distinct clans. Our objective was to determine whether the domains within a metacluster, annotated by one clan, are structurally related to those annotated by the other clan. To assess this, we calculated the TM-score for each pair of domains, comparing domains from the first Pfam clan with those from the second, then averaged the scores across all pairs (see Methods).

The results are presented in 4(b), where we analyzed Pfam labels with a minimum overlap of either 30% (violet curve) or 80% (red curve) with a corresponding DPCstruct domain. At the 30% minimum overlap, most metaclusters containing different Pfam labels do not suggest a structural relationship between the clans. This is likely because the Pfam labels are targeting different regions of the DPCstruct domains, rather than the same structural feature. However, when we increase the minimum overlap threshold to 80%, the TM-scores increase in value, reflecting a stronger structural alignment between the Pfam labels.

From this analysis, we identified hundreds of structurally related Pfam clans and families. While a comprehensive analysis of all cases is beyond the scope of this paper, we have included a list of all Pfam relationships in our dataset, which can be accessed online, in the hope that the community can make use of them. In the following section, we will discuss a few examples in detail to illustrate the type of relationships present in DPCstruct’s classification.

### Uncovering structural relationships among Pfam clans and families

The analysis of metaclusters annotated with different Pfam labels revealed numerous structural connections between Pfam families and clans. Here, we examine a few examples of both known and novel structural relationships, illustrating the type of connections present in our classification and how it can complement sequence-based annotations.

Figure 5(a) shows the superposition of two domains within MC28435: the orange domain from the cysteine-rich secretory or CAP family (PF00188) and the blue domain from the Eimeria tenella SAG family (PF11054). Despite their low sequence similarity, DPCstruct classifies the two domains as belonging to the same metacluster. Notably, the same relationship was also observed in a recent study on the structure of the SAG family [26]. In fact, based on this result, the recently released Pfam v37.0, included the SAG family as part of the CAP clan (CL0659), in agreement with our classification.

**Fig. 5:**
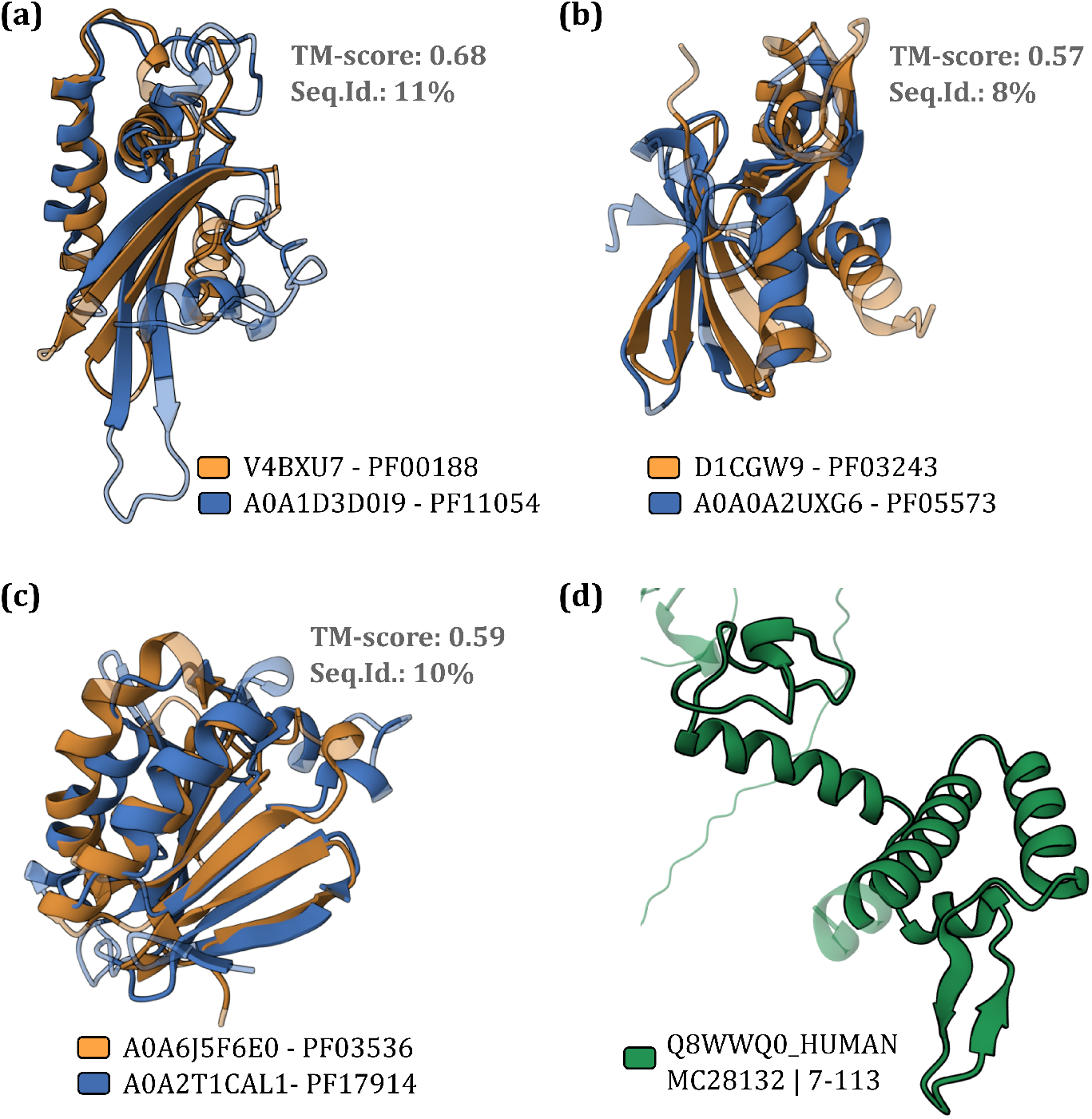
DPCstruct classification can capture structural relationships between distant Pfam families or clans and uncover putative novel structural families. (a)-(c) Examples of Pfam families relationships captured by a DPCstruct metacluster. For each example, we show the superposition of two domains members of a metacluster, each annotated by a different Pfam family. Superposition was calculated on the full DPCstruct domains using the RCSB PDB alignment tool (https://www.rcsb.org/alignment) [35, 36]. Transparency differentiates structurally aligned regions from non-aligned regions. (d) Partial view of the human protein Q8WWQ0, a pleckstrin homology domain-interacting protein, highlighting a putative new domain identified by DPCstruct as member of metacluster MC28132. This domain has no matching label in known domain databases (source: InterPro [7]).

Another example of DPCstruct’s ability to capture putative remote homologs is metacluster MC40874, which contains domains annotated by Pfam families MerB (PF03243) and NosL (PF05573), as shown in Figure 5(b). NosL, a copper chaperone, is part of the TRASH clan and is characterized by domains with well-conserved cysteine residues involved in metal coordination. MerB, on the other hand, plays a crucial role in bacterial detoxification pathways by cleaving Hg-carbon bonds. Although not part of the same Pfam clan, the two families are known to be structural homologs [27]. Furthermore, studies have suggested an evolutionary relationship between these families, supported by a possible link in their metal-binding sites [28, 29]. To further substantiate these claims, we performed an HHpred [30] comparison between the families, which yielded an 85% probability of homology.

Finally, we analyze metacluster MC46234, which contains domains from PF03536 and PF17914 (see Figure 5(d)). PF03536 contains phosphothreonine lyases that are used by pathogenic bacteria such as Salmonella to inactivate mitogen-activated protein kinases (MAPKs) in the host [31, 32]. PF17914 encompasses the HopA1 effector protein from Pseudomonas syringae, a bacterial pathogen that targets a wide range of species by interfering with the EDS1 complex [33]. A previous study suggested a weak structural relationship between these families [34]. DPCstruct, however, assigns them to the same metacluster, considering them structurally related. Furthermore, we compared these families using HHpred, which assigned a probability of homology of 90%. Both families involve effectors that interact with specific host proteins, albeit through different mechanisms. Despite these differences, the structural similarity identified by DPCstruct, together with the high probability of homology indicated by HHpred, suggests that they are evolutionary related. While further experimental validation is required, the example serves as a compelling use case of DPCstruct’s capability to uncover meaningful structural relationships that may not be immediately evident through sequence-based methods alone.

### The unknown metaclusters

Among the 28,246 metaclusters identified by DPCstruct, 24% exhibit no structural similarity to known protein families or folds. These “unknown metaclusters” represent putative novel protein domains that have yet to be characterized. To identify them, we conducted an extensive search of their representatives against the PDB, CATH, and Pfam databases, excluding all metaclusters containing any partial hits to these databases.

Unknown metaclusters are generally small in size, as larger metaclusters are more likely to be annotated. However, 91% have more than 5 elements with the largest metacluster containing 438 elements. To assess the quality of the structures these metaclusters are made of, we compared their TM-score, LDDT, and pLDDT distributions to those of metaclusters with known annotation. The distributions are indistinguishable, as shown in Figure 2(c) for pLDDT, indicating that unknown metaclusters are of the same structural quality as annotated ones and can be used as seeds for novel folds or superfamilies.

To illustrate how DPCstruct can be used as a seed to discover novel families or folds, we identified 10 unknown metaclusters, each containing at least one domain within a protein from the human proteome. A search for proteins containing these domains in the UniProt [37] and InterPro [7] databases provided further information about their function and existing domain annotations (see Supplementary Table 1). Most of these unknown domains are found in proteins with either full annotations, such as those provided by the PANTHER protein and gene classification database[38], or domain annotations in regions not overlapping DPCstruct labels. Notably, two of the selected unknown domains match Pfam-N domains, an extension to Pfam that uses deep neural networks to identify additional protein domains [39].

As an example from this list, we identified a previously unknown domain in the pleckstrin homology domain-interacting protein (PHIP). This protein, which is 1,821 amino acids long, is believed to be involved in the regulation of the insulin and insulin-like growth factor signaling pathways [40]. Although PHIP contains multiple known domains, no domains have been reported in the region shown in Figure 5(d), which DPCstruct identifies as part of metacluster MC28132.

## Discussion

Deep learning models have revolutionized the field of structural biology by generating hundreds of millions of predicted protein structures. While these models have proven to be an invaluable resource, taking full advantage of such a vast amount of data remains a critical challenge, which calls for the development of automatic labeling and annotation tools. In response to this need, we propose *DPCstruct*, a novel pipeline to automatically predict and group protein domains based on their structural similarity. The outcome is a classification of the protein space into structurally related domains, which can be used to build functional hypotheses.

Our work was made possible by recent advances, including freely available structural prediction databases such as AlphaFoldDB v4.0 and the development of open-source, fast, and accurate structural aligners like Foldseek [1, 2, 15]. In line with these principles of openness and accessibility, we release DPCstruct along with its associated database, with the hope that the community can use the pipeline as a stand-alone tool and the dataset as a complementary resource to enhance existing annotations.

Using DPCstruct on the Foldseek Clusters database we identified 28,246 domain clusters. These clusters are consistent with Pfam clan-level classification and further enrich it by using structural information to capture distant homologs. They also closely correspond to existing structure-based protein domain databases such as SCOP and CATH, identifying 88% and 95% of the folds present in each database respectively. More interestingly, 24% of the metaclusters do not match any of the aforementioned databases, representing a pool of putative novel folds, some of which are quite abundant in nature. This novel set of protein domain clusters offers a valuable resource for the community, providing potential new targets for functional studies.

Our classification also allows to address broader questions about the processes governing protein evolution. For example, we observed that the distribution of domains follows a power-law, even when considering proteins from different genomes, in agreement with protein evolution models typically applied to single genomes. Another related aspect is the extent to which proteins are successfully clustered. The Foldseek Cluster database contains a non-redundant set of 15 million proteins representing the full structural diversity of the AlphaFold database. The initial step of the pipeline, involving an all-vs-all local alignment, identifies only 16% of the input proteins with the minimum number of 10 hits required to be processed by DPCstruct. Among these, 53% are classified into metaclusters. The low level of local structural similarity between representative proteins can be attributed to their structural uniqueness or poor characterization by AlphaFold, as evidenced in the Supplementary Figure 1. Despite this, DPCstruct is able to identify almost all known folds, as the representatives that it clusters are the most abundant in nature, representing roughly 65% of all proteins in the AlphaFoldDB. The number increases to 80% if one consider partial hits to our metaclusters.

Compared to sequences, the clustering of protein structures presents new challenges, mainly due to the increased complexity and degrees of freedom in structural data. One significant issue is the occurrence of inconsistent alignments, where a structure may align well with two other structures individually, but those two structures do not align well with each other. Such instances complicates the clustering process and is more prevalent in structural data than in sequence-based clustering. To mitigate this problem, we have taken a conservative approach by imposing stricter thresholds in the all-versus-all alignments, thereby also limiting the sensitivity of our method. A simple solution would be to relax the Foldseek search thresholds, especially the threshold on the E-value. However, this adjustment would increase the number of false positives in our primary clusters, introducing errors into our results. To address this, we plan to modify the clustering criteria to better handle noise. One possible solution is to use the Advanced Density Peaks algorithm instead of the standard algorithm [41]. In addition, the modular and fast implementation of DPCstruct will allow us to update and improve our classification as new and improved versions of structural aligners or prediction tools are released. Future efforts will also include the integration of sequence information into the clustering procedure, along with improvements in our distance estimation using protein representations based on deep learning models.

In this study, we focused on the AlphaFoldDB, which covers almost all UniProtKB entries, with the exception of viral and long proteins. We showed that DPCstruct retrieves the vast majority of known information from this well-studied database, that it unveils some previously unknown relations among protein families, and that it detects more than 6,000 putative novel structural domains. Building upon these findings, we believe DPCstruct could be effectively applied to the expanding field of metagenomics, for instance on the ESMAtlas database [20], which includes a broader range of structures beyond those in UniProtKB, potentially revealing further novel domains.

## Methods

### Dataset

The dataset used comprises roughly 15 million proteins, representatives of the Foldseek Clusters database, a full-protein clustered version of the AlphaFold database, which contains over 200 million proteins (see Figure 1(a)). This clustering was performed at a 50% sequence identity and 90% sequence overlap threshold, followed by a 90% structure identity clustering, as presented by Barrio-Hernandez et al. [16]. PDBs together with their respective per-residue pLDDT were obtained using Google Cloud Services, gs://public-datasets-deepmind-alphafold-v4. Foldseek v7 createdb module was used for building the input dataset.

### Pipeline overview

For this study, we present DPCstruct, an enhanced version of the DPCfam pipeline, which succeeds the previous iteration used in our previous work for clustering protein sequence-based databases, including UniRef50 [6] and UHGP [19], to identify protein domains. The DPCstruct pipeline completed its analysis in under 48 hours on a 4-node cluster, utilizing 120 cores of Intel Xeon Gold 6126 processors. What follows is an outline of the pipeline, highlighting the specific modifications implemented for this study. For a more comprehensive description of the methodology, please refer to Russo et al. [6].

The pipeline takes as input the representative proteins of the Foldseek Clusters database, described in the previous section. By using a non-redundant version of the AlphaFoldDB, we avoid the overrepresentation of proteins which could introduce biases in our clustering algorithm. In this respect Foldseek clustering as performed in Ref. [16] can be seen as the first essential step in our pipeline.

We then generate pairwise local alignments for all pairs of proteins using Foldseek v7 [15], a new structural alignment tool. Foldseek uses a structural alphabet to transform the structural alignment problem into a sequence-based one. This adaptation allows the use of well-established and optimized tool, MMseqs2 [42], to efficiently solve the alignment problem. For the all-versus-all alignment step we used the default settings of the software, with a sensitivity value set to 7.5 and an E-value threshold of 0.001, a rather restrictive value chosen to reduce the occurrence of random alignments.

Given that we are dealing with protein structure predictions, an additional filtering step was necessary based on the average predicted LDDT value reported by AlphaFold [2]. Alignments where either the query or the target sequence region had an average residue pLDDT smaller than or equal to 60% were discarded. This threshold selection aligns with AlphaFold guidelines, which established a lower bound for the per-residue pLDDT between 50% and 70%.

After the all versus all local alignment, we perform two clustering steps that are the core of the DPCstruct algorithm. The first step, called primary clustering, takes all local alignments associated with a single query protein, assesses the density of the alignments in the query space, and clusters them accordingly using density peak clustering [43]. Alignments in regions of low density are filtered out, while alignments around density peaks are merged. This process is performed independently for each query protein.

The second step consists of grabbing all clusters identified for all proteins and clustering them together to generate a set of metaclusters. The procedure is based once again on the Density Peak Clustering algorithm, but this time performed on an abstract space of primary clusters, where the distance is calculated as the number of common elements between clusters normalized by the size of the smaller cluster. The final output is a set of putative protein families we call metaclusters. Each consists of a list of structurally related protein domains.

### Primary cluster selection criteria

The primary modification of the algorithm, aside from replacing the local alignment tool to accommodate protein structures and incorporating an ad-hoc filter to manage structure predictions, involved a change in the primary cluster selection criteria. This adjustment significantly enhanced the algorithm’s ability to identify primary clusters. For the Foldseek Cluster representatives database, this resulted in at least a doubling of the number of primary clusters identified.

The Density Peaks Clustering algorithm, introduced by Rodriguez and Laio [43], identifies the peaks of a density distribution by utilizing a decision graph that plots the density, *ρ*, against the minimum distance to a point with a higher density, *δ*. Outliers in this plot represents the density peaks of the distribution. In the context of primary clusters, the objective is to identify regions within a protein with a high number of overlapping alignments. In particular, the distance between alignments is defined as one minus the ratio of the intersection to the union of the segments, and *ρ* is the number of overlapping alignments. Initially, these outliers were identified as the top-scoring data points ranked based on the product of *ρ* and *δ*. In the current version, we employ a double threshold, set as *δ* ≥ 0.4 and *ρ* ≥ 10. This means we identify clusters containing at least 10 alignments, with the center of the cluster having a distance of at least 0.4 to another alignment with higher density.

### Consistency score

For the comparison between DPCstruct and Pfam, we used Pfam 36.0 seed alignments restricted to the representative proteins of the Foldseek Cluster dataset. To measure the agreement between the classifications, we used a consistency score introduced in Barrio-Hernandez et al.[16], which extends the normalized mutual information score to accommodate our case, where a domain may have multiple labels from the same classification.

The consistency score between a metacluster and Pfam classification is determined by considering the subset of domains within the metacluster that overlap with any specific Pfam label. For each pair of proteins within this subset, we count the number of common Pfam labels and divide it by the total number of labels in the reference protein. This quantity is not symmetric, as labels in one protein may contain, but not necessarily be equal to, the labels of the other protein. The overall consistency score is then calculated by averaging over all pairs of proteins.

### Comparison of Pfam labels within a metacluster

For a metacluster with two different Pfam labels, we separated its domains based on their Pfam annotation. We then computed the pairwise TM-score of all members in one group against the members in the other group. The alignment was performed considering only the region of the DPCstruct domain overlapping with the respective Pfam label. By averaging these values, we obtained a TM-score per metacluster.

## Data availability

The dataset associated with this study, including the DPCstruct classification and the data needed to reproduce all plots, can be found at the following Zenodo repository: https://zenodo.org/doi/10.5281/zenodo.13334295.

## Code availability

Plots were generated using Python 3.12.0, Matplotlib 3.8.0, Pandas 2.1.1 and RCSB custom Mol* viewer (https://www.rcsb.org/)[36][35]. The DPCstruct algorithm is available as free and open-source software (GPL-3.0 license) at https://github.com/RitAreaSciencePark/DPCstruct.

## Acknowledgments

The authors warmly thank Marco Celoria for collaboration in the initial stages of this project. The authors acknowledge the AREA Science Park supercomputing platform ORFEO made available for conducting the research reported in this paper and the technical support of the Laboratory of Data Engineering staff. A.A., and A.C. were supported by the European Union – NextGenerationEU within the project PNRR “PRP@CERIC” IR0000028 - Mission 4 Component 2 Investment 3.1 Action 3.1.1.

## Supplementary Material

**Supplementary Tab. 1:** List of unknown metaclusters whose representative domains, selected as the cluster centers, belongs to the human proteome. Descriptions and annotations were obtained from UniProt [37] and InterPro [7] databases. The column of additional annotations only considers Pfam or PANTHER labels.

| Metacluster | Metacluster size | Representative Protein | Domain Range | Protein description | Additional annotations |
| --- | --- | --- | --- | --- | --- |
| MC45663 | 25 | B2RCN4 | 23-264 | Transcription regulation | PTHR16088 |
| MC28132 | 24 | Q8WWQ0 | 7-113 | PH-interacting protein | PF00400, PF00439, PTHR16266 |
| MC4520 | 18 | Q9UKK3 | 1554-1714 | Mono-ADP-ribosyltransferase | PF00533, PF00092, PF00644, PF08487, PTHR46530 |
| MC5376 | 17 | Q8NEX5 | 1-80 | Serine-type endopeptidase inhibitor activity | - |
| MC11593 | 14 | Q9UG01 | 1662-1748 | Intraflagellar transport protein | PF00400, PTHR15722 |
| MC30005 | 14 | Q0P6D6 | 782-936 | Coiled-coil domain-containing protein | PTHR14817 |
| MC55004 | 8 | Q8N2J1 | 163-268 | mRNA splicing | PF07842, PTHR12214 |
| MC31975 | 6 | B3KTF3 | 1-107 | Ataxin-7-like transcription regulator | PF16135, PTHR15117 |
| MC14068 | 6 | A0A3B3ISK8 | 1-47 | Zinc finger protein | PTHR23057 |
| MC44855 | 2 | A0A804HKJ4 | 9-77 | Involved in glycosylation | PF13439, PTHR13036 |

**Supplementary Fig. 1:**
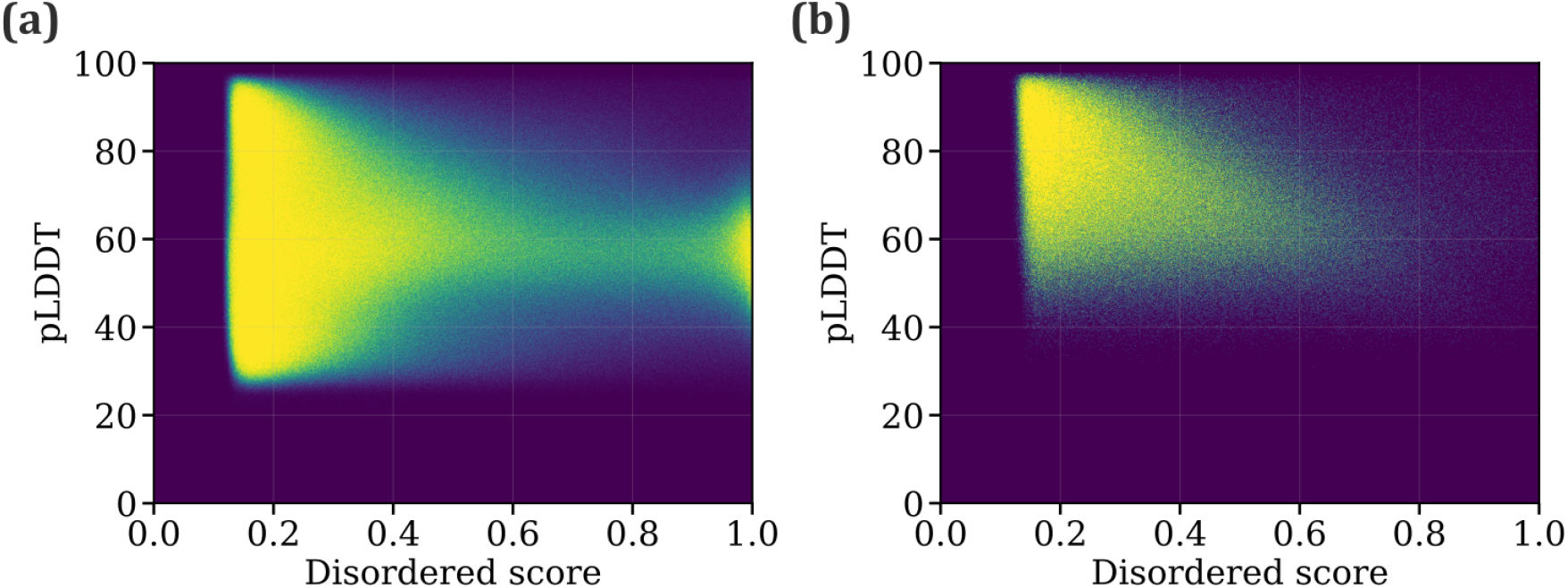
Relationship between mean pLDDT as reported by AlphaFold [1] and mean predicted intrinsically unstructured score calculated with AIUPred [44] for (a) all Foldseek Cluster representatives [16] excluding fragments (15.3 million proteins) and (b) the subset of representatives labeled by DPCstruct. The means were calculated over the amino acids. Only 31% of all the proteins have a pLDDT > 60 and a disordered score < 0.4. Most of DPCstruct-labeled proteins are found in this region. Color scale is proportional to the density of proteins in that region, with yellow representing the highest density.

